# Effects of junk-food on food-motivated behavior and NAc glutamate plasticity; insights into the mechanism of NAc calcium-permeable AMPA receptor recruitment

**DOI:** 10.1101/2023.05.16.540977

**Authors:** Tracy L. Fetterly, Amanda M. Catalfio, Carrie R. Ferrario

## Abstract

In rats, eating obesogenic diets increase calcium-permeable AMPA receptor (CP-AMPAR) transmission in the nucleus accumbens (NAc) core, and enhances food-motivated behavior. Interestingly these diet-induced alterations in NAc transmission are pronounced in obesity-prone (OP) rats and absent in obesity-resistant (OR) populations. However, effects of diet manipulation on food motivation, and the mechanisms underlying NAc plasticity in OPs is unknown. Using male selectively-bred OP and OR rats, we assessed food-motivated behavior following ad lib access to chow (CH), junk-food (JF), or 10d of JF followed by a return to chow diet (JF-Dep). Behavioral tests included conditioned reinforcement, instrumental responding, and free consumption. Additionally, optogenetic, chemogenetic, and pharmacological approaches were used to examine NAc CP-AMPAR recruitment following diet manipulation and ex vivo treatment of brain slices. Motivation for food was greater in OP than OR rats, as expected. However, JF-Dep only produced enhancements in food-seeking in OP groups, while continuous JF access reduced food-seeking in both OPs and ORs. Reducing excitatory transmission in the NAc was sufficient to recruit CP-AMPARs to synapses in OPs, but not ORs. In OPs, JF-induced increases in CP-AMPARs occurred in mPFC-, but not BLA-to-NAc inputs. Diet differentially affects behavioral and neural plasticity in obesity susceptible populations. We also identify conditions for acute recruitment of NAc CP-AMPARs; these results suggest that synaptic scaling mechanisms contribute to NAc CP-AMPAR recruitment. Overall, this work improves our understanding of how sugary, fatty food consumption interacts with obesity susceptibility to influence food-motivated behavior. It also extends our fundamental understanding of NAc CP-AMPAR recruitment; this has important implications for motivation in the context of obesity as well as drug addiction.

## I. INTRODUCTION

As obesity rates continue to rise globally (Swinburn et al., 2011), it is increasingly important to understand the neural and behavioral consequences of over-consuming calorie-dense palatable foods. In humans, consumption of obesogenic diets (i.e., calorie dense, high-fat/high-sugar) alter the function of brain reward centers including the Nucleus Accumbens (NAc; Small, 2009; Horstmann et al., 2011; Stice et al., 2013; Volkow et al., 2013). Furthermore, greater activations in the NAc in response to food cues are observed in obesity-susceptible people before weigh gain (Demos et al., 2012; Murdaugh et al., 2012). These data highlight interactions between diet and obesity-susceptibility that contribute to over-eating and weight gain. However, our understanding of neurobehavioral differences in obesity-prone and -resistant populations is limited, and relatively little is known about how obesogenic diets affect NAc function and food-seeking behaviors.

In rodents, consuming obesogenic diets produces a number of alterations in neuronal function (Ferrario et al., 2016), including enhancements in NAc dendritic spine density and excitatory transmission (Tukey et al., 2013; Counotte et al., 2014; Dingess et al., 2017), and decreases in dendritic spine density in the prefrontal cortex (PFC; Dingess et al., 2017). Furthermore, obesity-susceptible rats show enhanced diet-induced NAc glutamatergic plasticity, including potentiation of NAc glutamate synapses (Brown et al., 2017) and increased calcium-permeable AMPA receptor (CP-AMPAR) transmission (Oginsky et al., 2016; Alonso-Caraballo et al., 2021; Nieto et al., 2023). This suggests that that food-seeking and feeding behaviors that rely on the NAc may also be altered by obesogenic diets.

Consistent with this, consumption of obesogenic foods enhances cue-triggered food-seeking (Dingess et al., 2017; Derman and Ferrario, 2018a). Activity of NAc CP-AMPARs is required for the expression of cue-triggered food-seeking (Crombag et al., 2008; Derman and Ferrario, 2018b) and increases in NAc CP-AMPAR expression are associated with enhancements in this behavior in obesity-prone, but not obesity-resistant rats. However, results describing how eating obesogenic foods alter the willingness to work for food are mixed, with both enhancements and reductions reported (Finger et al., 2012; Matikainen-Ankney et al., 2023). Thus, there is a need to further tease apart factors that influence food motivated behavior in states of over-abundance and obesity. In addition, despite established differences in NAc function and incentive-motivation in obesity-prone vs -resistant populations, how obesity susceptibility interacts with diet to alter food motivation and NAc glutamatergic plasticity is unclear.

Here we examined how eating a sugary, fatty “junk-food” (JF) diet affects several key aspects of food-seeking and feeding behavior. This was done both while rats were maintained on this diet, and after a brief JF “deprivation” period (JF-Dep) during which rats only have access to ad lib standard lab chow. Additionally, to further our understanding of the mechanisms underlying CP-AMPAR recruitment during JF-Dep, we investigated the conditions that permit increases in NAc CP-AMPAR transmission. Given that the medial PFC (mPFC) and basolateral amygdala (BLA) send direct glutamatergic projections to the NAc and are both involved in food-seeking behavior (Cardinal et al., 2002; Holland and Petrovich, 2005; Christoffel et al., 2021), we also determined the input specificity of CP-AMPAR upregulation.

## II. METHODS

### Animals

Male obesity-prone (OP) and obesity-resistant (OR) rats bred in house (50-60 days old at the start of each study) were used (Levin et al., 1997). Rats were singly housed for behavior experiments and pair-housed for electrophysiology experiments (reverse 12-h light/dark cycle). Procedures were approved by The University of Michigan Committee on the Use and Care of Animals in accordance with AAALAC guidelines.

### Food and Weight

Rats were weighed daily unless otherwise specified and home cage food intake was measured throughout. Chow (CH) controls were maintained on standard lab chow (Lab Diet 5L0D: 4 kcal/g; 13.6% fat, 28.9% protein, 57.5% carbohydrates; % of caloric content), while experimental junk-food (JF) groups were given free access to JF made in house (Oginsky et al., 2016; Robinson et al., 2015; 4.5 kcal/g; 19.6% fat, 14% protein, and 48% carbohydrates). This JF diet was a mash of Ruffle potato chips (40g), Chips Ahoy (130g), Nesquik (130g), Jiff peanut butter (130g), powdered Lab Diet 5L0D (200g) and 180mL of water. JF Deprivation (JF-Dep) consisted of removing JF from the home cage and returning rats to free-access standard lab chow.

### Behavior

All behavioral testing was conducted in standard operant boxes within sound attenuating chambers (Med Associates; one session/day). A general description of each experiment is given first followed by details of each behavioral approach.

### Experiment 1

Effects of junk-food and junk-food deprivation in OP rats. OP rats were food restricted (85-90% free-feeding bodyweight) and given 12 Pavlovian Conditioning sessions. Rats were then assigned to chow, JF and JF-Dep groups counterbalanced by behavior during the final three conditioning sessions; they had ad lib access to food for the remainder of the study. Next, rats underwent a single conditioned reinforcement test session followed by instrumental training and testing. Finally, rats underwent free-consumption testing (see timeline Fig. 1A).

**FIG. 1:**
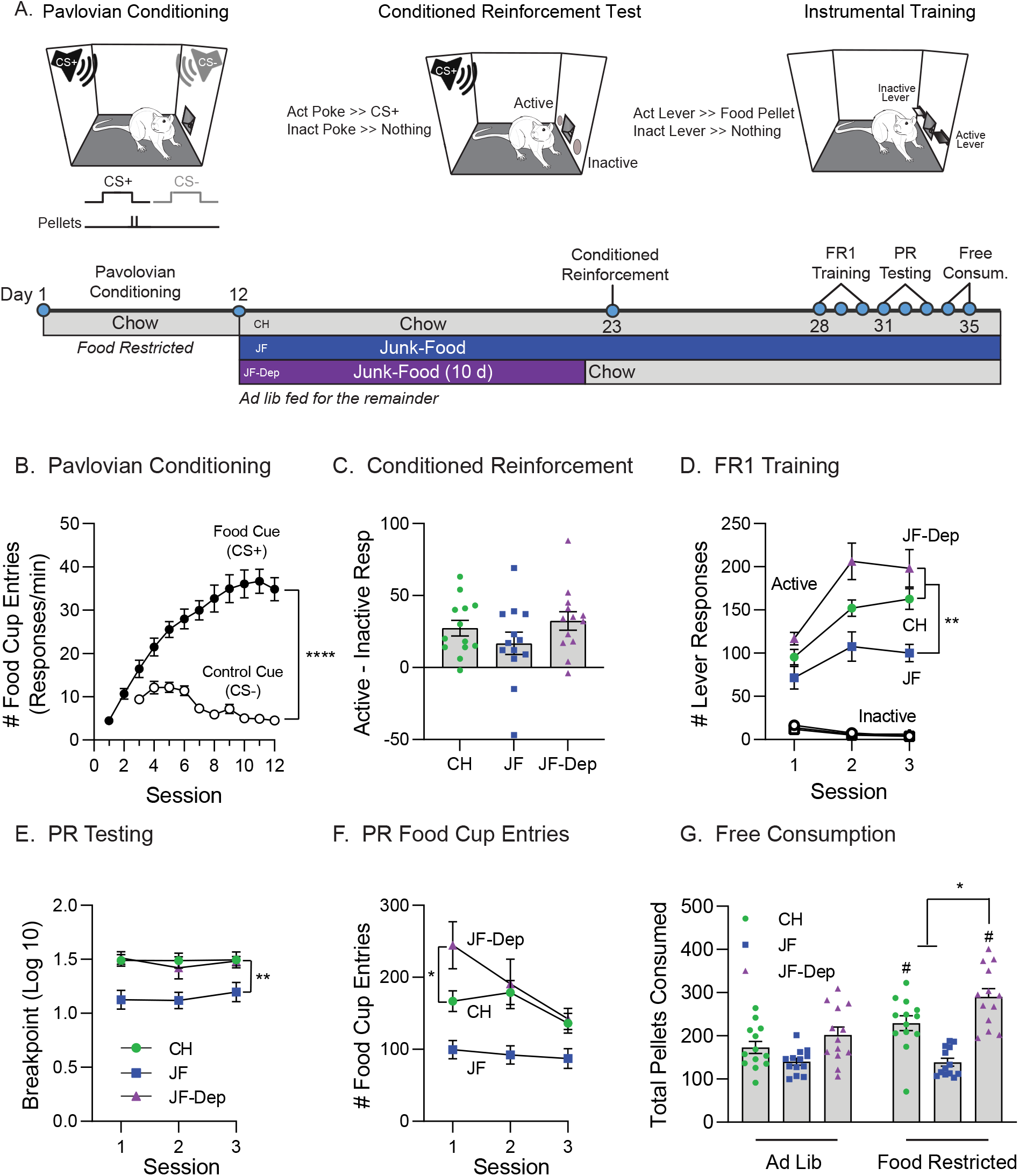
Effects of junk-food and junk-food deprivation on behavior in obesity-prone rats. A)Cartoons of behavioral conditioning, testing and experimental timeline. Obesity-prone rats (N=39) first received Pavlovian conditioning and were then divided into chow (CH), junk-food (JF), and junk-food deprivation (JF-Dep) groups (N=13 rats/group) before undergoing behavioral testing. B) Pavlovian conditioning. All rats learned to discriminate between the Food Cue (CS+) and Control Cue (CS-; ^****^p<0.0001, main effect of cue). C) The magnitude of conditioned reinforcement (active responses - inactive responses) did not differ between groups. D) Number of lever responses during each of three fixed ratio 1 (FR1) training sessions. All rats preferentially responded on the active vs inactive lever. Rats consuming JF showed lower active lever responding compared to rats that underwent JF-Dep or those that remained on CH (^**^p<0.01, Sidak’s post-test). E) Breakpoint during each of three progressive ratio (PR) tests was similar across repeated testing. Break point was reduced in JF vs CH and JF-Dep groups, but similar between CH and JF-Dep groups (^**^p<0.01, Sidak’s post-test). F) Number of food cup entries during each PR test. Food cup entries were elevated during the first session in the JF-Dep group compared to CH (*p<0.05, Sidak’s post-test). G) Pellets consumed during free consumption testing conducted when rats were fed ad lib or food restricted (4.5h) prior to the test. Pellet consumption did not differ across diet groups when rats were tested in the ad lib state (left bars). Thus, lower responding during FR1 and PR sessions in the JF group is not likely due to reductions in value of the food pellets per se. Food restriction increased pellet consumption in CH and JF-Dep groups (#p<0.05, Sidak’s post-test ad lib vs. restricted), but had no effect on consumption in the JF group. Furthermore, following food restriction the JF-Dep group consumed more pellets than either CH or JF groups (^*^p<0.05, Sidak’s post-test). All data are presented as mean *±* SEM

### Experiment 2

Effects of junk-food and junk-food deprivation in OP vs OR rats. OP and OR rats were assigned to diet groups counterbalanced by starting weight within strain and had ad lib access to food throughout. Following diet manipulation, rats underwent instrumental training and testing (see timeline Fig. 2A).

**FIG. 2:**
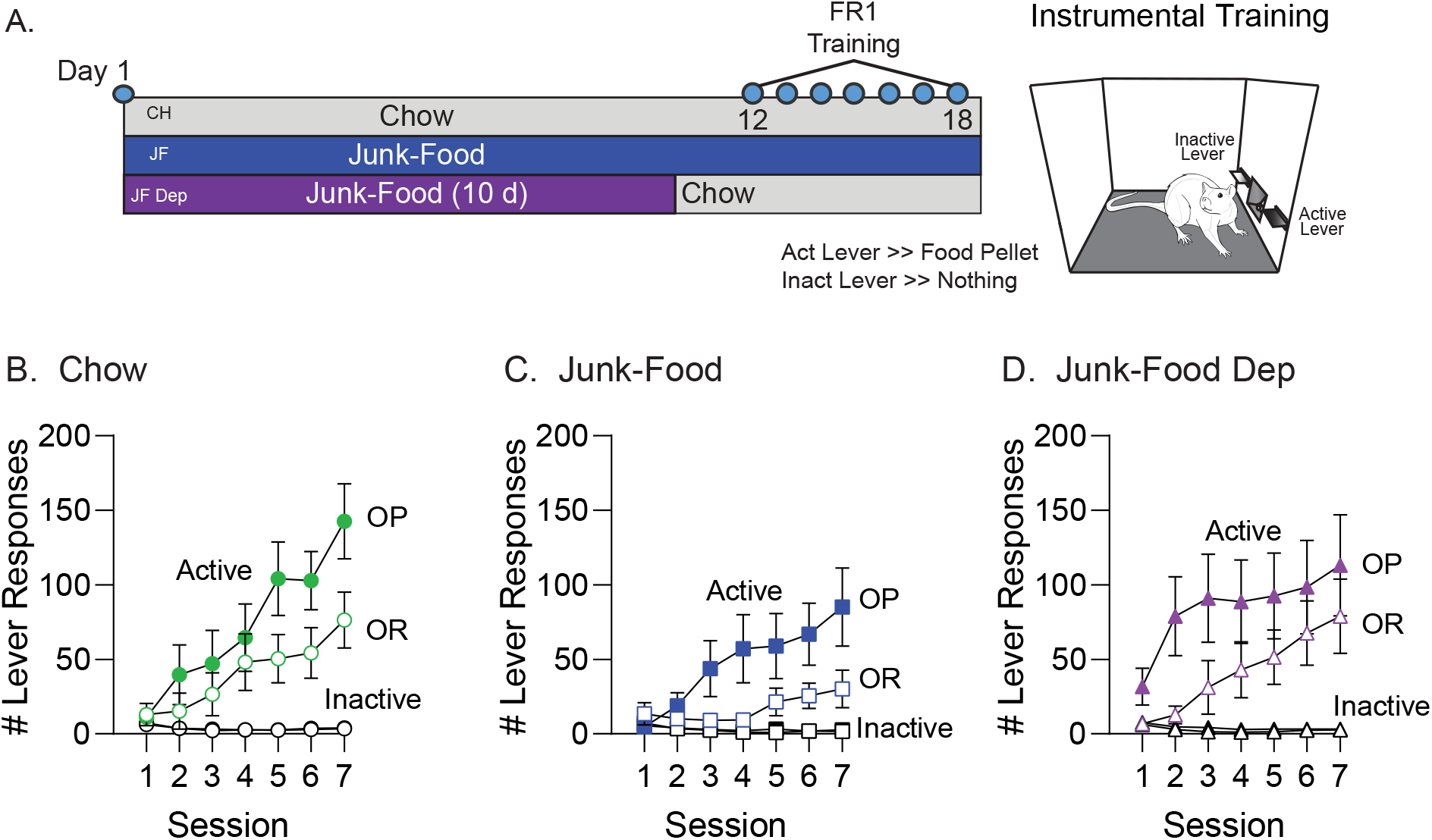
Effects of junk-food and junk-food deprivation on instrumental responding in obesity-prone and obesity-resistant rats. A) Cartoon of instrumental training, testing, and experimental timeline. Obesity-prone (OP) and obesity-resistant (OR) rats were first assigned to chow (CH), junk-food (JF), and junk-food deprivation (JF-Dep) groups (N=12/group) before undergoing instrumental training (FR1). B-D) Number of active and inactive lever responses during each training session organized by diet group. All groups responded preferentially on the active lever, and inactive lever responses remained low and stable across sessions. For all conditions, OP groups showed greater active lever responses than OR groups. In addition, free access to JF in the home cage reduced active lever responses in both OPs and ORs compared to their chow controls. JF-Dep (D) resulted in enhanced active lever responses early in training (sessions 1-4) in OP rats as compared to OR rats (see results for stats).

### Pavlovian Conditioning

Procedures were adapted from (Holland et al., 2002 and Petrovich et al. 2002). A white nose and a tone were used, counter-balanced across CS+/CS-assignment. Rats first underwent two sessions where presentation of one cue (CS+, 10s, 8 presentations/session) was followed by delivery of two food pellets (45 mg, Bioserv, #F0059, banana; 6.3% fat, 20.2% protein, 52% carbohydrates; % of caloric content). Next rats were given ten discrimination sessions where presentation of a second cue (CS-, 10s) that was never paired with food pellets was interspersed with CS+ presentations (4 CS+, 4 CS-, 8 presentations total; inter-trial-interval [ITI] 2-6 min, 32 min/session). Entries into the food cup during CS presentations and the ITI were recorded throughout.

### Conditioned Reinforcement Test

Conditioned reinforcement for the CS+ was assessed in a single session (30 min, Robinson et al., 2015). Responses in one nosepoke port (active) resulted in CS+ presentation (3s), but no food pellets. Responses in the other port (inactive) produced no outcome. Active and inactive responses were recorded throughout.

### Instrumental Training and Testing

Rats were trained to press one lever (active) to obtain the same food pellet used above (fixed ratio 1; FR1, 30 min/session). A second lever (inactive) was present but had no consequences. Rats had three FR1 training sessions followed by three progressive ratio (PR) testing sessions. During each PR session the number of active lever presses required to obtain each subsequent food pellet delivery increased exponentially (5e(delivery#X0.2)-5), adapted from Richardson and Roberts (1996). The PR session ended when rats did not meet the next ratio requirement within 30 min (i.e., breakpoint).

### Free Pellet Consumption

Rats underwent two free pellet consumption tests (30 min/session). During each test, rats were placed in the operant box and allowed to freely consume food pellets that had been placed in the food cup. The first test occurred under ad lib fed conditions, whereas the second occurred following a 4.5h home-cage food restriction period (no food in cage). The total number of pellets consumed during each session was measured.

### Electrophysiology

Surgeries and Viral Microinjection: Rats were anesthetized with isoflurane and injected intracranially with AAV constructs expressing either Chronos or hM4D(Gi)-DREADD (500 nL/hemisphere; 1*μ*L/min). The analgesic carprofen (2.5 mg/kg, i.p.) was administered at the time of surgery and once every 24h for 48h following surgery. For optical stimulation experiments, pAAV-Syn-Chronos-GFP (Addgene viral prep 59170-AAV1; http://n2t.net/addgene:59170; RRID:Addgene 59170; gifted by E. Boyden) was infused into either the mPFC or BLA (mPFC: AP +3.00, ML +/- 0.60, DV -3.0; BLA: AP -2.28, ML +/- 5.00, DV -7.2). For DREADD experiments, pAAV-Syn-hM4D(Gi)-mCherry (Addgene viral prep 50475-AAV8; http://n2t.net/addgene:50475; RRID:Addgene 50475; gifted by B. Roth) was infused into the mPFC. Controls for DREADD studies were injected with either the pAAV-Syn-Chronos-GFP or pAAV-Syn-mCherry. Electrophysiological recordings were made from these rats 3-4 weeks after surgery.

### Whole-Cell, Voltage-Clamp Recordings

All reagents were purchased from Millipore Sigma unless otherwise noted. Brain slices containing the NAc (bregma +0.96-2.52) were prepared as previously described (Oginsky et al., 2016; Fetterly et al. 2021). Briefly, coronal sections (300 *μ*) were prepared using a vibratome (Lecia Biosystems). Slices recovered in oxygenated artificial cerebrospinal fluid (aCSF; in mM: 122.5 NaCl, 25 NaHCO3, 12.5 Glucose, 1 NaH2PO4, 1 L-ascorbic acid, 2.5 KCl, 2.5 CaCl2, 1 MgCl2; 295-305 mOsm, pH 7.45) at 37ºC for 30 min.

Established whole-cell patch-clamping approaches were used (Oginsky et al., 2016, Alonso-Caraballo et al., 2021). All recordings were made using Clampex 10.7-11.1 and analyzed using Clampfit 10.7-11.1 (Molecular Devices). Patch pipettes were pulled from 1.5 mm borosilicate glass capillaries (WPI; 3-7 MΩ resistance) and filled with a solution containing (in mM): 140 CsCl, 10 HEPES, 2 MgCL2, 5 Na+-ATP, 0.6, Na+-GTP, 2 QX314; 285 mOsm; pH 7.3. Whole-cell voltage-clamp recordings of AMPA receptor-mediated EPSCs were made at -70 mV in the presence of picrotoxin (50 *μ*M). Electrically evoked EPSCs (eEP-SCs) were elicited by local stimulation (0.05 to 0.30 mA square pulses, 0.1 ms, delivered every 20s) using a bipolar electrode placed 300 *μ*m lateral to recorded neurons. Optically evoked EPSCs (oEPSCs) were elicited under the control of a CoolLED pE-300ultra and passed through a LED-FITC-A-OMF filter cube to produce blue wavelength light (0.2 ms, delivered every 20s). Epifluorescent images were used to confirm injection sites prior to recording (Olympus, BX43 with X-Cite LED). The minimum amount of current or light intensity needed to elicit a synaptic response with <15% variability in amplitude was used. Cells for which access resistance changed by >20% across the recording were discarded. EPSCs were recorded before and after application of the CP-AMPAR selective antagonist Naspm (200*μ*M; as in Conrad et al., 2018; Oginsky et al., 2016) to measure CP-AMPAR mediated transmission, or Clozapine-N-Oxide (CNO; 10*μ*M; as in Fetterly et al., 2019) to activate DREADDs. For ex vivo studies of CP-AMPAR recruitment, slices were incubated in oxygenated aCSF containing either CNO (10*μ*M) or the NMDAR antagonist (2R)-amino-5-phosphonovaleric acid (APV; 50*μ*M) for at least two hours before recordings began. All stock drug solutions were diluted in ddH2O, except for picrotoxin and CNO (diluted in DMSO).

### Statistical Analysis

One-tailed or two-tailed t-tests, one-way, two-way, or three-way ANOVAs were used. Sidak’s multiple comparisons were used for post-hoc analysis (GraphPad Prism 9). Comparisons were made between OP, OR and diet groups within each experiment. All Ns are given in the figure captions. Data in all figures are shown as average *±* SEM.

## III. RESULTS

### A. Effects of junk-food and junk-food deprivation in OP rats

After Pavlovian conditioning rats were separated into chow (CH), junk-food (JF), and JF-deprivation (JF-Dep) groups (Figure 1A). Rats discriminated between the CS+ and CS-, making more entries into the food cup during CS+ vs CS-presentations (Figure 1B; main effect of cue, *F*_(1,76)_=102.4, p<0.0001; session *X* cue interaction, *F*_(9,684)_=25.06, p<0.0001). Rats were then given a single conditioned reinforcement test after 10 days of JF, or 10 days of JF plus 24h of JF-Dep; controls remained on chow throughout (Figure 1A). All groups showed conditioned reinforcement, responding more in the active vs inactive port to receive a presentation of the CS+ (data not shown: main effect of port, *F*_(1,72)_=38.72, p<0.0001). To simplify comparisons, the magnitude of conditioned reinforcement (active responses - inactive responses) was compared across groups (Figure 1C). All groups showed similar motivation to work for presentations of the CS+ alone (no effect of diet group, *F*_(2,36)_=1.46, p=0.24).

We next evaluated instrumental responding in these same rats (Figure 1D). During training, all groups responded preferentially on the active vs inactive lever and inactive responses were low and stable throughout (Figure 1D; main effect of lever, CH: *F*_(1,24)_=247.9, p<0.0001, JF: *F*_(1,24)_=83.67, p<0.0001, JF-Dep: *F*_(1,24)_=111.3, p<0.0001). In addition, the magnitude of active responding differed across groups, with JF rats responding the least and JF-Dep responding the most across all three sessions (Figure 1D; active responses: main effect of diet group, *F*_(2,36)_=12.44, p<0.0001). Motivation to work for the food pellets was assessed in three PR testing sessions. Behavior was stable across these tests (Figure 1E; no effect of session, *F*_(1.724,62.06)_=0.99, p=0.36). Similar to FR1 training, breakpoint was reduced in JF vs both CH and JF-Dep groups (Figure 1E; main effect of diet group, *F*_(2,36)_=8.57, p=0.0009; no group *X* session interaction, *F*_(4,72)_=0.59, p=0.67). Interestingly, during the first PR session the number of food cup entries was greater in the JF-Dep group compared to the CH group (Figure 1F; main effect of diet group, *F*_(2,36)_=10.95, p=0.0002; post-test: CH vs JF-Dep, p=0.05), even though breakpoints were similar across these groups. Thus, rats in the JF-Dep group show greater instrumental responding during FR1 testing, and increased food-seeking (i.e., food cup entries) during the first PR session.

Figure 1G shows pellet consumption during free access testing when rats were fed ad lib or following a 4.5hr food restriction period. Pellet consumption did not robustly differ across diet groups when rats were tested in the ad lib state. Thus, lower responding during FR1 and PR sessions in the JF group is not likely due to reductions in value of the food pellets per se. Food restriction increased pellet consumption in CH and JF-Dep groups (Figure 1G; main effect of state, *F*_(1,36)_=32.53, p<0.0001; diet group x state interaction, *F*_(2,36)_=9.78, p=0.0004; post-test ad lib vs. restricted: CH, p=0.0013; JF-Dep, p<0.0001). However, no such effect of food restriction was found in the JF group (post-test ad lib vs fasted: p=0.99). In addition, the JF-Dep group consumed significantly more pellets than either the CH or JF-Dep group when tested following food restriction (post-test: CH vs JF-Dep, p=0.015; JF vs JF-Dep, p<0.0001). These results are consistent with greater instrumental responding in the JF-Dep group when effort is relatively low (Figure 1D) and with enhanced food cup entries during the first PR test (Figure 1F).

### B. Effects of junk-food and junk-food deprivation in obesity-prone vs obesity-resistant rats

We next compared effects of JF and JF-Dep on instrumental responding in OP and OR rats. After two days of JF-Dep, all rats began instrumental training (Figure 2A). All groups responded preferentially on the active vs inactive lever and inactive responses were low and stable throughout (Figure 2B-D; main effect of lever, CH: *F*_(1,22)_=25.42, p<0.0001, JF: *F*_(1,22)_=13.44, p=0.0014, JF-Dep: *F*_(1,22)_=16.65, p=0.0005). The OP-CH group made more active responses than OR-CH (Figure 2B; session *X* strain interaction, *F*_(6,132)_=2.43, p=0.029). Although active responding was generally lower in JF groups, the pattern of greater responding in OP vs OR groups persisted (Figure 2C: session *X* strain interaction, *F*_(6,132)_=3.03, p=0.0083). This pattern was disrupted by JF-Dep, with increased active responding in earlier sessions, and less separation between OPs and ORs (Figure 2D).

Within session active responses during the initial acquisition period for OR (A-D) and OP (E-H) groups are shown in Figure 3. For OR rats, elevations in active responding begin to emerge by the fourth training session, with visually greater responding in OR-CH and OR-JF-Dep groups as compared to OR-JF (Figure 3D). Active responding in OR-CH and OR-JF-Dep groups was similar and remained fairly stable throughout the session (Figure 3D, Session 4: no time *X* diet interaction, *F*_(10,165)_=1.18, p=0.31). In contrast, increased responding in the OP-JF-Dep group was present even during the first session compared to OP-CH and OP-JF groups (Figure 3E: main effect of diet F_(2,33)_=3.45, p=0.044, main effect of time, *F*_(2.757,90.98)_=3.26, p=0.028, time *X* diet interaction, *F*_(10,165)_=2.22, p=0.019). This same general pattern is seen in sessions two and three (Figure 3F,G). In addition, active responding in the OP-JF-Dep group was greater early vs late within training sessions two through four. Indeed, by session four, responding is fairly consistent across time in OP-CH and OP-JF groups (similar to the OR rats), while responding in the OP-JF-Dep group was significantly greater early in the session, and declined across time (Figure 3H: main effect of time, *F*_(2.08,68.65)_=7.63, p=0.00009, time *X* diet interaction, *F*_(10,165)_=2.33, p=0.013). Thus overall, OPs with a history of JF-Dep showed greater active lever responding than OP-CH and OP-JF groups, an effect that was not seen in the ORs.

**FIG. 3:**
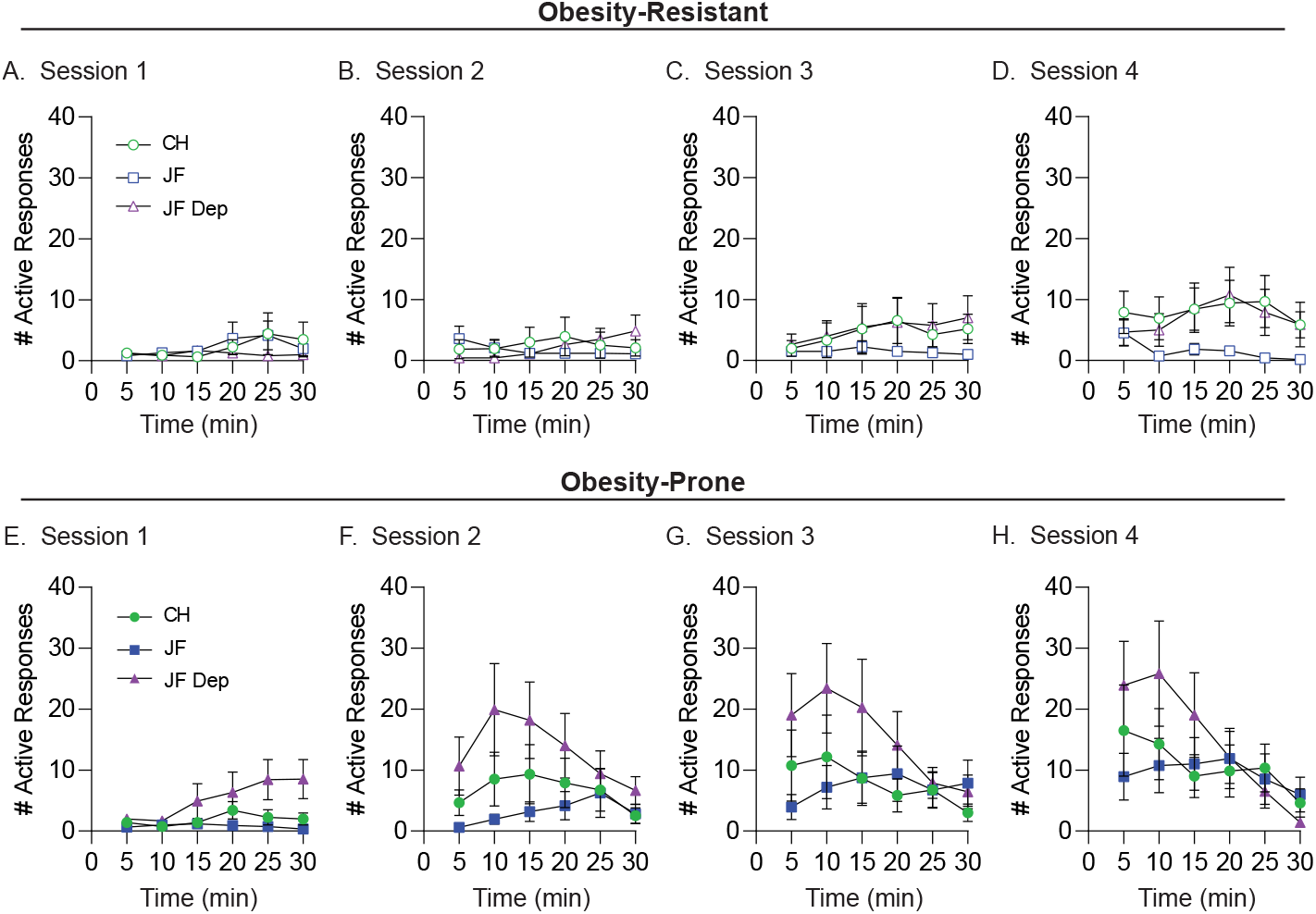
Within session active lever responding across the first four FR1 training sessions. A-D) Number of active lever responses per five minutes during training sessions 1-4 in OR groups. Active responding was low and similar in CH and JF-Dep groups across these sessions, and begins to visually increase compared to the JF group by session 4 (D). E-H) Number of active lever responses per five minutes during training sessions 1-4 in OP groups. In contrast to all other groups, increases in active lever responses emerged earlier during training in the JF-Dep group. In addition, there was a shift in activity, such that by the third and fourth sessions (G,H) responding was highest during the first half of the session and then declined across time in the OP-JF-Dep group. This is contrasted by a slower emergence of increased responding across sessions and stable responding through the entire 30min session in OP-CH and OP-JF groups.

Figure 4 shows food intake (A,B) and weight gain (C,D) for rats in Experiment 1 (upper graphs) and Experiment 2 (lower graphs). The dotted line indicates when rats in the JF-Dep group were placed back on ad lib chow. For both experiments, rats given free access to JF consumed more food than those given free access to chow (Figure 4A: main effect of diet, *F*_(2,36)_=6.81; Figure 4B: main effect of diet, OR: *F*_(2,33)_=13.50, p<0.0001, OP: *F*_(2,33)_=16.53, p<0.0001). In addition, there were no differences in food intake between OP-JF and OR-JF groups across 10 days of JF exposure (Figure 4B: no effect of strain, CH: *F*_(1,22)_=0.51, p=0.48, JF: *F*_(1,22)_=0.13, p=0.72, JF-Dep: *F*_(1,22)_=0.086, p=0.77), and food intake dropped briefly when rats in the JF groups were placed back on ad lib chow. In regard to weight gain, for both experiments, rats in the JF group gained significantly more weight than those in the chow groups (Figure 4C: main effect of diet group, *F*_(2,36)_=6.65, p=0.0035; diet group *X* time interaction, *F*_(46,828)_=10.17, p<0.0001; Figure 4D: main effect of diet group, OR: *F*_(2,33)_=10.70, p=0.003, OP: *F*_(2,33)_=16.65, p<0.0001; main effect of time, OR: *F*_(1.378,45.46)_=810.2, p<0.0001, OP: *F*_(1.435,47.37)_=937.0, p<0.0001; diet group *X* time interaction, OR: *F*_(36,594)_=7.61, p<0.0001, OP: *F*_(36,594)_=13.06, p<0.0001). As expected, weight gain in the OP-JF-Dep group was similar to OP-CH controls by the end of the study (Figure 4C: Sidak’s post-test; Day 35, CH vs. JF-Dep, p=0.13), whereas the OP-JF group gained more weight than OP-CH (**=Sidak’s post-test, Day 35, CH vs JF, p=0.0013). Finally, in Experiment 2, OP rats gained more weight than OR rats, regardless of diet group (Figure 4D: main effect of diet, CH: *F*_(1,22)_=13.00, p=0.0016, JF: *F*_(1,22)_=12.56, p=0.0018, JF-Dep: *F*_(1,22)_=6.02, p=0.023; Sidak’s post-test Day 18: OP-CH vs OP-JF: p<0.0001), although the OR-JF group did gain more weight than the OR-CH group (Sidak’s post-test Day 18: OR-CH vs OR-JF p=0.0024). Overall, differences in food-intake and weight gain did not correspond to behavioral differences between experimental groups and chow controls, or between OP and OR rats.

**FIG. 4:**
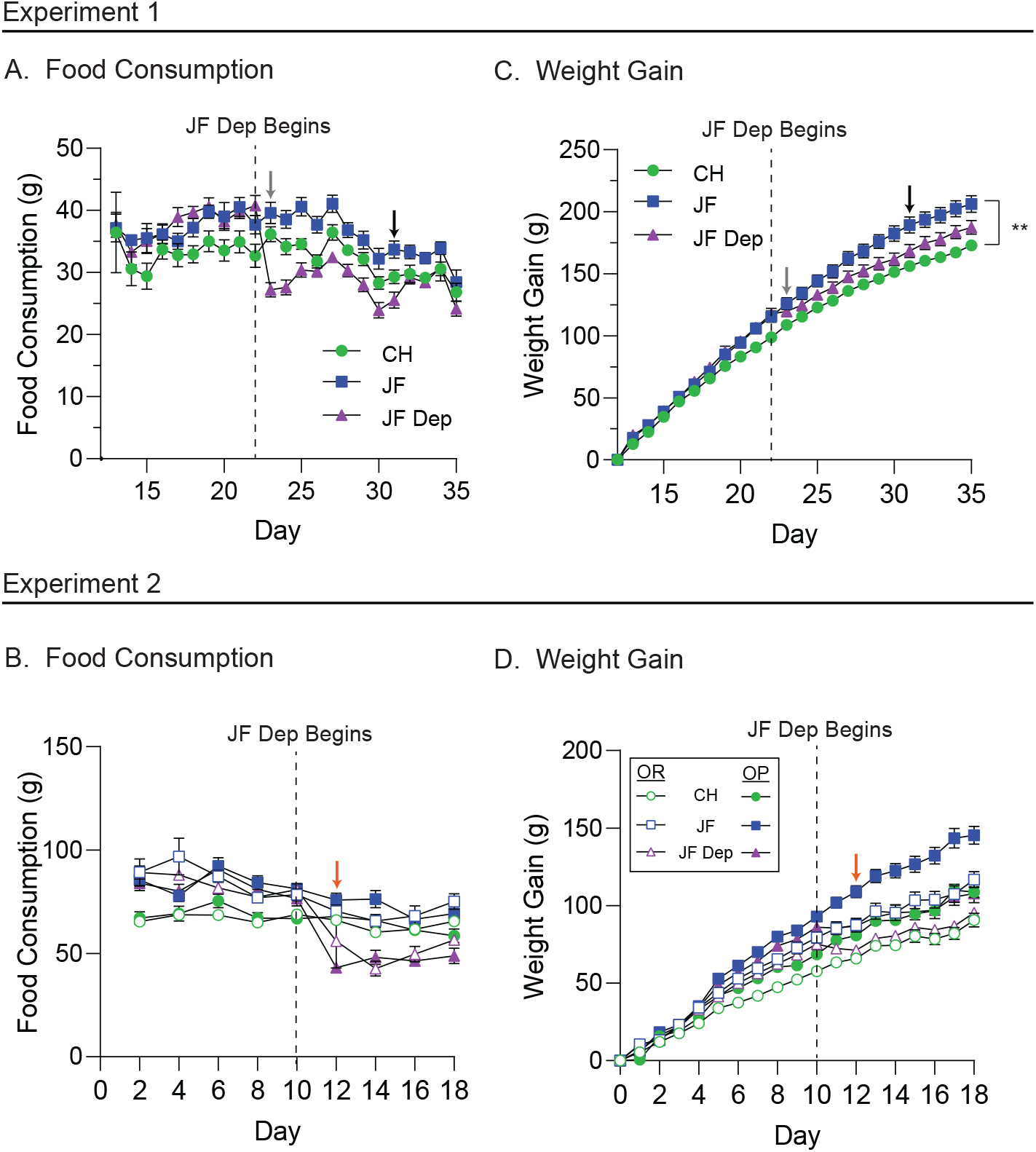
Weight gain and food consumption corresponding to behavioral experiments. A-B) Food consumption for rats in Experiment 1 (N=13 rats/group) and Experiment 2 (N=12 rats/group), respectively. For both experiments, rats given free access to JF consumed more food than those given free access to chow. C-D) Weight gain of rats in Experiment 1 and 2, respectively. For both experiments, rats consuming JF gained significantly more weight than those on chow (^**^p<0.01, Sidak’s post-test). In Experiment 2, regardless of diet group, OP rats gained more weight than OR rats. The dashed line in each panel indicates when the JF-Dep group was returned to ad lib chow. Arrows indicate the conditioned reinforcement test (blue), the first day of progressive ratio testing (black), and the first day of instrumental training (orange). Overall, although expected differences in food intake and weight gain were observed between OP and OR rats, differences in food-motivated behavior between these lines and in response to diet manipulation did not correspond to differences in food intake or weight gain.

### C. Recruitment of NAc CP-AMPARs and input specificity

The same JF-Dep manipulation used above results in increases in NAc CP-AMPAR transmission in OP but not OR rats, and a JF deprivation period is necessary for this increase (Oginsky et al., 2016; Alonso-Caraballo et al., 2021). Synaptic recruitment of CP-AMPARs can be induced by prolonged reductions in excitatory transmission via synaptic scaling mechanisms in other neuronal systems (Turrigiano et al., 1998; Ju et al., 2004; Sutton et al., 2006). Therefore, we next determined whether incubating slices containing the NAc in the NMDAR antagonist APV was sufficient to recruit CP-AMPARs to synapses (Figure 5A). In slices from OP rats, APV incubation increased the sensitivity to Naspm in both chow and JF groups (Figure 5B-C: main effect of slice treatment, *F*_(1,38)_=11.76, p=0.0015; no main effect of group, *F*_(1,38)_=0.33, p=0.57; no group *X* treatment interaction, *F*_(1,38)_=0.00024, p=0.99). In contrast, APV incubation of slices from ORs did not alter Naspm sensitivity (Figure 5D-E: t_(13)_=0.26, p=0.79). Thus, dampening excitatory transmission was sufficient to increase NAc synaptic CP-AMPAR transmission in OP but not in OR rats; this effect was similar in slices from OP-CH and OP-JF groups.

**FIG. 5:**
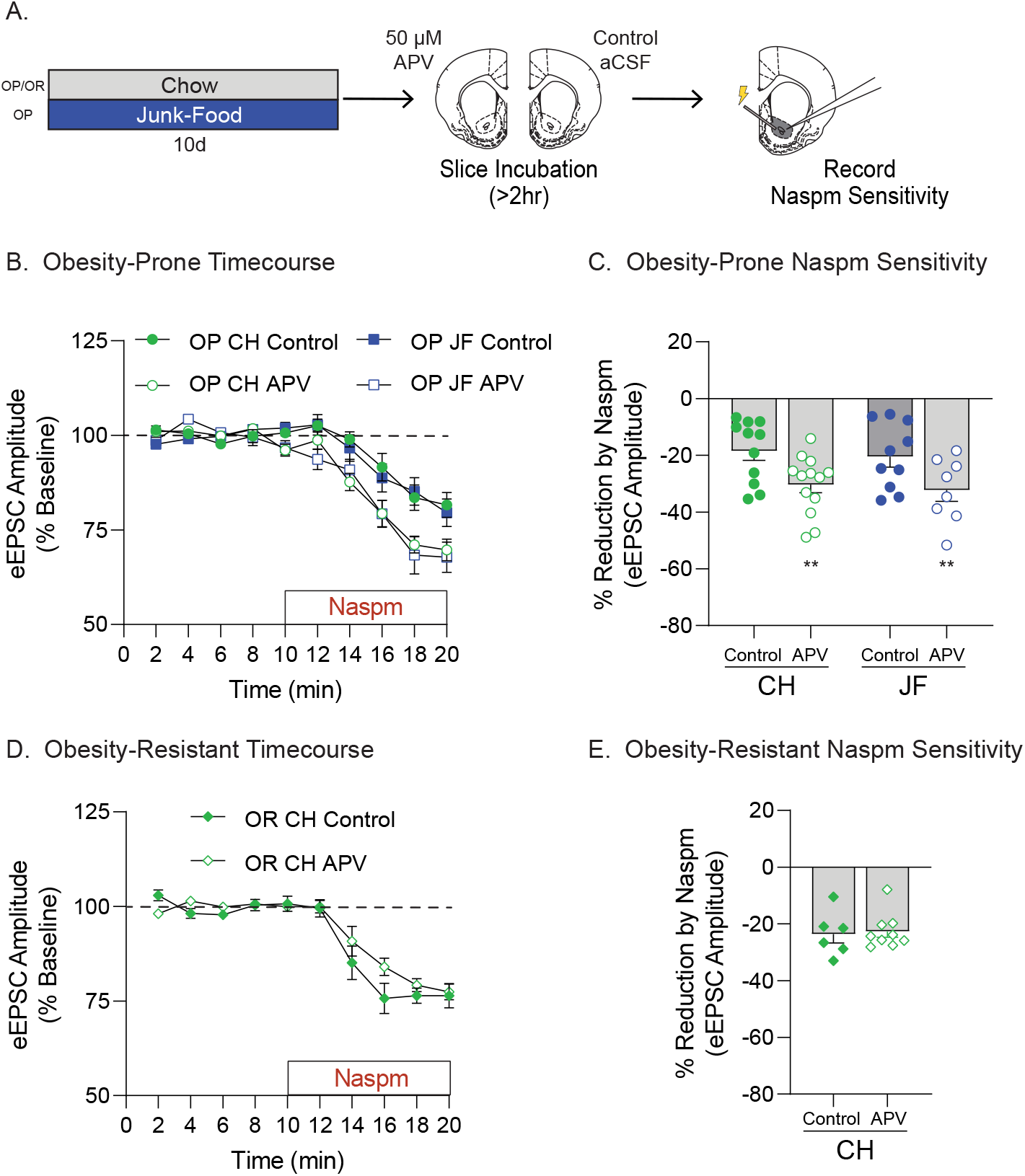
Ex vivo NMDA receptor blockade is sufficient to recruit NAc synaptic CP-AMPARs in obesity-prone, but not obesity-resistant rats. A) Schematic of experimental timeline. After 10 days of free access to junk-food (JF) or chow (CH), slices containing the NAc were prepared for electrophysiology. All slices were immediately incubated in either aCSF (Control) or aCSF with APV (50 *μ*M) for at least 2 hours prior to recording. B) Timecourse showing effects of bath application of the CP-AMPAR antagonist Naspm (200 *μ*M) on eEPSC amplitude in recordings made from obesity-prone (OP) groups (N = number of rats, number of cells; CH: Control N=10,1 APV N=11,13; JF: Control N=9,10 APV N=8,8). C) Percent reduction in eEPSC amplitude following Naspm application (avg. of last two minutes of drug wash-on from B). APV incubation increased Naspm sensitivity (i.e., CP-AMPAR-mediated transmission) in OP-CH and OP-JF groups (^**^p<0.01, main effect of slice treatment). D) Timecourse showing effects of bath application of Naspm on oEPSC amplitude in recordings made from obesity-resistant (OR) groups (CH: Control N=6,6 APV N=9,9). E) Percent reduction in eEPSC amplitude following Naspm application (avg. of last two minutes of drug wash-on from D). APV incubation did not alter CP-AMPAR-mediated transmission in ORs.

We next identified which synaptic inputs CP-AMPAR transmission is enhanced in by comparing the effects of JF-Dep in mPFC or BLA inputs to the NAc using optogenetics combined with Naspm sensitivity (Figure 6A). Figure 6A shows an example of viral expression within mPFC and BLA. Figure 6B,C show data from oEPSCs in mPFC-to-NAc inputs. Naspm sensitivity of mPFC-to-NAc inputs was increased only in the OP-JF-Dep group (Figure 6B-C: main effect of strain, *F*_(1,32)_=8.64, p=0.0061; OP-CH vs OP-JF-Dep: t_(22)_=2.25, p=0.017, OR-CH vs OR-JF-Dep: t_(10)_=0.61, p=0.28, based on a priori planned comparisons). Figure 5D,E show data from oEPSCs in BLA-to-NAc inputs. Interestingly, no effects of JF-Dep were found in BLA-to-NAc inputs (Figure 6E: no effect of strain, *F*_(1,24)_=0.012, p=0.91, no effect of diet group, *F*_(1,24)_=0.31, p=0.58, no group *X* strain interaction, *F*_(1,24)_=1.30, p=0.27).

**FIG. 6:**
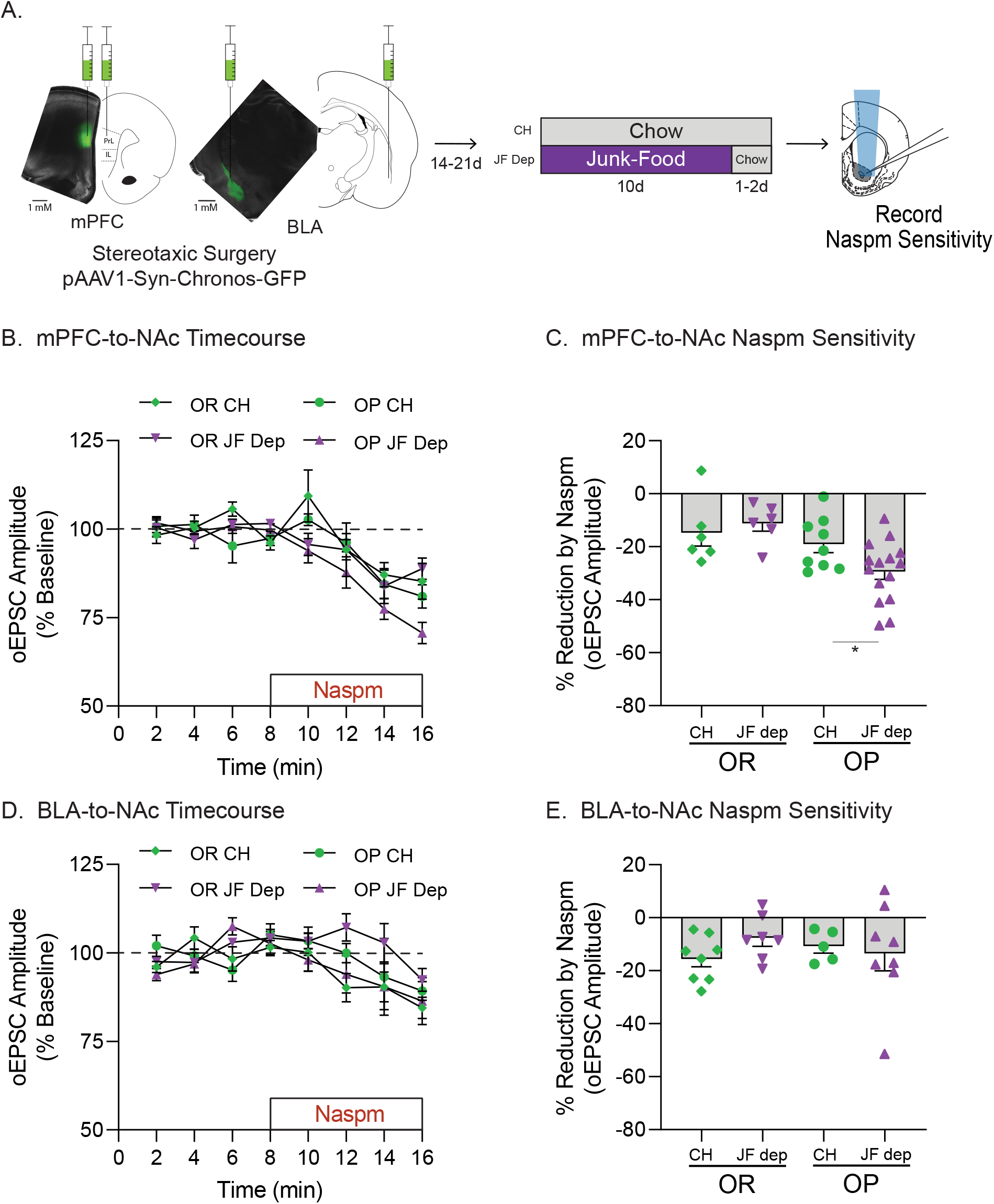
Junk-food deprivation increases CP-AMPAR transmission in mPFC-to-NAc, but not BLA-to-NAc, synapses. A) Schematic of experimental timeline. pAAV-Syn-Chronos-GFP was bilaterally infused into either the mPFC or BLA of obesity-prone (OP) and obesity-resistant (OR) rats. Rats were then given 2-3 weeks to recover and allow for viral expression before diet manipulation (Chow or JF-Dep). Slices were then prepared for electrophysiology using optical stimulation of mPFC or BLA inputs to the NAc core. B) Timecourse showing effects of bath application of the CP-AMPAR antagonist Naspm (200 μM) on oEPSC amplitude in mPFC-to-NAc synapses (OR: CH N=5,6 JF-Dep N=5,6; OP: CH N=6,9 JF-Dep OP N=9,15). C) Percent reduction in oEPSC amplitude following Naspm application (avg. of last two minutes of drug wash-on from B). JF-Dep increased Naspm sensitivity in mPFC-to-NAc synapses of OPs but not ORs (^*^p<0.05, OP-CH vs OP-JF-Dep: one-tailed t-test based on a priori planned comparisons). D) Timecourse showing effects of bath application of Naspm on oEPSC amplitude in BLA-to-NAc inputs (OR: CH N=4,8 JF-Dep OR N=4,7; OP: CH N=4,5 JF-Dep N=7,8). E) Percent reduction in oEPSC amplitude following Naspm application in BLA-to-NAc inputs (avg. of last two minutes of drug wash-on from D). JF-Dep did not alter CP-AMPAR transmission in BLA- to-NAc inputs of OPs or ORs.

Given that CP-AMPAR increases occur in mPFC-to-NAc synapses following JF-Dep in OPs, and that reducing excitatory transmission was sufficient to increase NAc CP-AMPAR transmission, we next determined whether reducing activity of mPFC terminals within the NAc was sufficient to recruit CP-AMPARs. This was accomplished by expressing a Gi-coupled DREADD (hM4D(Gi)) in the mPFC and bath applying the agonist CNO to coronal slices containing the NAc and mPFC terminals. Figure 7A shows an example of viral expression within mPFC. Slices from one hemisphere were incubated in CNO while slices from the opposite hemisphere of the same rat were incubated in vehicle aCSF. This allowed for comparison of CNO effects within the same set of slices. CNO incubation increased Naspm sensitivity of the eEPSC amplitude compared to vehicle aCSF (Figure 7B: *t*_(5)_=4.14, p=0.0045), suggesting that reducing activity of mPFC terminals within the NAc is sufficient to increase synaptic CP-AMPAR transmission. To verify that CNO does not alter eEPSC amplitude on its own, and that activating hM4D(Gi) receptors on mPFC terminals in NAc slices decreases synaptic activity, control recordings in slices with and without hM4D(Gi)-expression were conducted. Acute CNO wash-on produced small, but significant reductions in eEPSC amplitude in slices expressing hM4D(Gi), but not in control slices (Figure 7C: main effect of time, *F*_(3.648,47.42)_=2.72, p=0.045, time *X* viral injection interaction, *F*_(9,117))_=2.43, p=0.014).

**FIG. 7:**
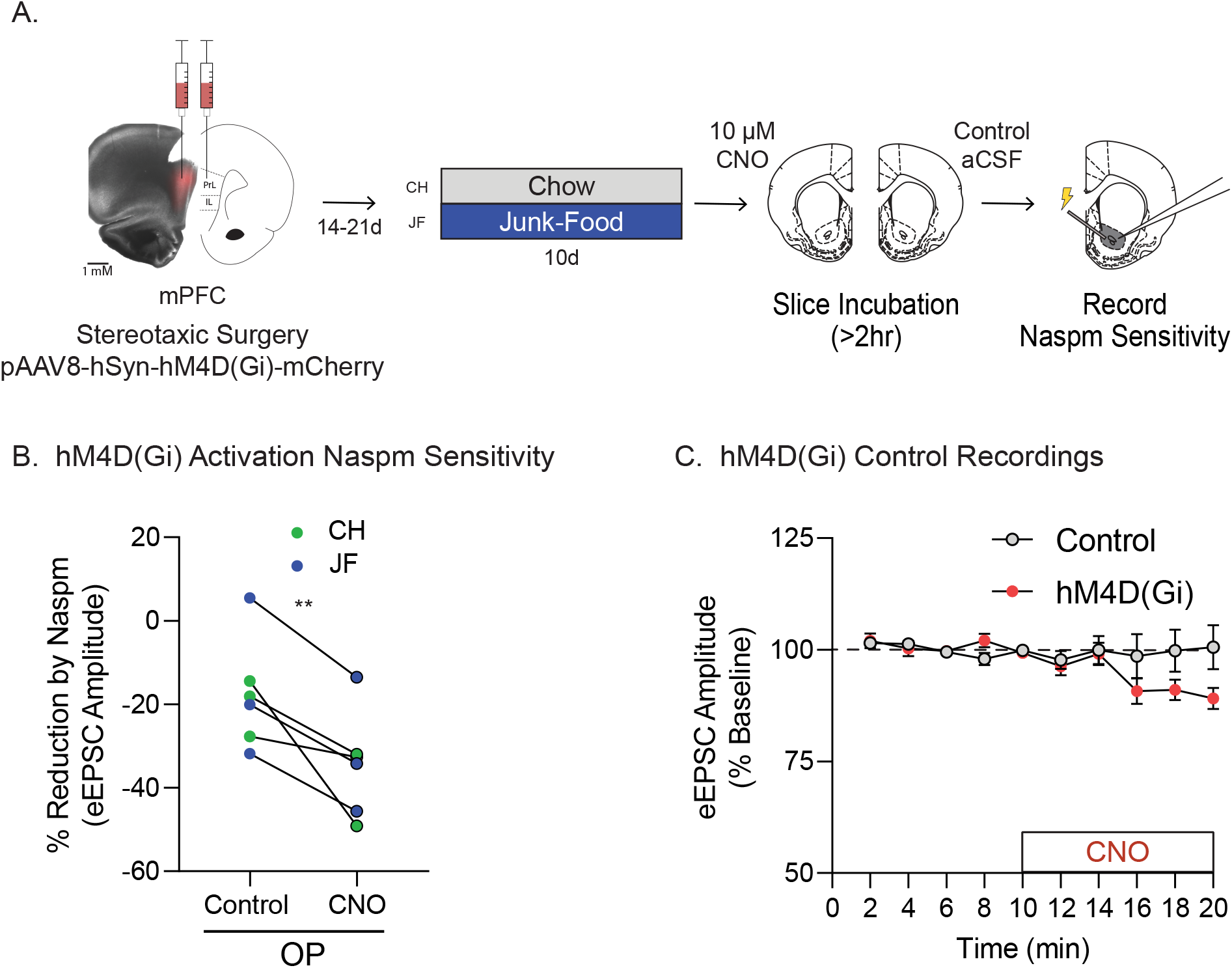
Reducing mPFC-to-NAc activity via activation of hM4D(Gi) receptors ex vivo is sufficient to recruit synaptic NAc CP-AMPARs in obesity-prone rats.A) Schematic of experimental timeline. pAAV-Syn-hM4D(Gi)-mCherry was bilateral infused into the mPFC of obesity-prone (OP) rats. Rats were then given 2-3 weeks to recover and allow for viral expression before starting the diet manipulation (Chow or JF). Slices containing the NAc were then prepared for electrophysiology. All slices were immediately incubated in either aCSF plus vehicle (0.1% DMSO; Control) or aCSF containing CNO (10 μM) for at least 2 hours prior to recording. B) Summary graph showing effect of CP-AMPAR antagonist Naspm (200 *μ*M) on eEPSC amplitude in cells from OP-CH (blue) and OP-JF (green) groups. Pairs of cells indicate recordings from slices obtained from the same rat (e.g., left hemisphere incubated in CNO, right incubated in control aCSF). CNO incubation significantly increased Naspm sensitivity in OP-CH and OP-JF groups (N=6,6/condition; ^**^p¡0.01, one-tailed t-test). C) Timecourse of control recordings in slices from additional chow-fed control rats with and without hM4D(Gi) expression. There was no effect of CNO on eEPSC amplitude when hM4D(Gi) was not expressed, while CNO decreased eEPSC amplitude when hM4D(Gi) was expressed in mPFC-to-NAc synapses (Control: N=6,6; hM4D(Gi): N=9,9).

## IV. DISCUSSION

We examined interactions between susceptibility to obesity and diet-induced neuronal and behavioral plasticity using established OP and OR models. Continuous junk-food access reduced food-seeking in both OPs and ORs, whereas consumption of junk-food followed by a period of deprivation (JF-Dep) enhanced food-seeking only in OPs. The persistence of behavioral effects beyond the removal of junk-food is consistent with long-lasting diet-induced neuroplasticity. Furthermore, we found that NAc CP-AMPAR increases induced by JF-Dep occur in mPFC, but not BLA, inputs to the NAc. Additionally, decreasing excitatory transmission by blocking NMDARs, or reducing activity of mPFC-to-NAc inputs ex vivo was sufficient to increase NAc CP-AMPAR transmission in OPs. These data highlight behavioral and physiological changes that may contribute to diet-induced obesity and provide insights into the mechanism of NAc CP-AMPAR recruitment.

### A. Effects of junk-food consumption on behavior

In OP rats, consumption of JF did not alter willingness to respond for the presentation of a food cue (Figure 1C) compared to CH controls. Yet in these same rats, instrumental responding for food pellets was reduced compared to CH controls (Figure1 D,E). This dichotomy of sustained responsivity to the food cue itself despite decreased motivation to work for food is consistent with results from outbred male rats where junk-food consumption enhances conditioned approach, without altering cue potentiated feeding (Derman and Ferrario, 2018a). Furthermore, junk-food-induced reductions in motivation for food in OPs was not related to lower motivational value of the food pellets per se, as rats in the junk-food group consumed similar amounts of these pellets as chow controls during free consumption testing (Figure 1G, ad lib). When OP and OR groups were studied side-by-side (Figure 2A), responding was generally greater in OPs than ORs, and was reduced by junk-food diet in both groups (Figure 2). Thus, despite general enhancements in motivation in OPs vs ORs, both groups were able to adjust their behavior in response to their dietary environment, working less for food pellets when JF was freely available in the home cage.

### B. Effects of junk-food deprivation on behavior

In Experiment 1, active lever responding was higher in OPs with a history of JF-Dep compared to OP-CH controls (Figure 1D). In addition, despite reaching equivalent breakpoints to the OP-CH group during PR testing, the OP-JF-Dep group had more entries into the food cup during the first PR session (Figure1F). This enhanced checking behavior is consistent with enhancements in motivation, although this difference did not persist during subsequent PR testing. Furthermore, when OP and ORs were tested side-by-side, we again found evidence for enhancements in motivation following JF-Dep only in OPs. Specifically, throughout FR1 training, active lever responding was indistinguishable between OR-JF-Dep and OR-CH controls (Figure 3A-D). In contrast, the OP-JF-Dep group showed elevated active lever responding during FR1 training that was largely due to the emergence of increased active responding earlier in training compared to all other groups (Figure 3E-H). Thus, when the overall effort to obtain food was relatively low, OPs that underwent JF-Dep showed more rapid acquisition of instrumental responding than their CH counterparts. Finally, when free pellet consumption was measured after food restriction, the OP-JF-Dep group also showed the greatest increase, consuming even more pellets than OPs maintained on chow (Figure 1G). This difference cannot be attributed to weight gain, as both the OP-CH and OP-JF-Dep groups gained a similar amount of weight across the study (Figure 4). In addition, these shifts in motivation and the regulation of food intake occurred when home cage food access during the deprivation period was unrestricted and was similar to food intake of chow controls (Figure 4). These aspects, in combination with effects of JF-Dep on neural function (discussed below), point to potential antecedents of obesity and have implications for the relatively poor long-term outcomes of current behavioral weight-loss strategies.

### C. Recruitment of NAc CP-AMPARs

CP-AMPARs have largely been studied in the context of drugs of abuse, fear conditioning, and neurodegeneration (Clem and Huganir, 2010; Wolf and Tseng, 2012; Dong et al., 2017; Guo and Ma, 2021), and have roles in both Hebbian and homeostatic plasticity (Man, 2011). For example, activity of NAc CP-AMPARs is required for cue-triggered food- and drug-seeking (Conrad et al., 2008; Derman and Ferrario, 2018b). Yet to date, mechanisms specific to the recruitment of NAc CP-AMPARs (which lack the GluA2 subunit) vs ‘traditional’ calcium-impermeable-AMPARs (CI-AMPARs), which contain the GluA2 subunit, have not been identified (Werner et al., 2017). Here, we found that decreasing synaptic activity via inhibition of NMDARs (*>*2h APV treatment) was sufficient to recruit CP-AMPARs in the NAc of OPs, but not ORs (Figure 5). This is reminiscent of homeostatic plasticity in cultured neurons that involves the “scaling up” of AMPARs in response to prolonged decreases in excitatory transmission (i.e., synaptic scaling), discussed further below.

With this in mind, we hypothesized that NAc core CP-AMPAR increases that occur following a necessary period of JF deprivation (Alonso-Caraballo et al., 2021) may result from reductions in excitatory input to the NAc. To pursue this idea, we first identified which synaptic inputs CP-AMPAR transmission is enhanced in using an optogenetic approach. We focused on mPFC-to-NAc and BLA-to-NAc inputs because these provide direct inputs to the NAc core and because these regions influence food-motivated behavior via interactions with the NAc (Cardinal et al., 2002; Holland and Petrovich, 2005; Christoffel et al., 2021). CP-AMPAR increases following JF-Dep were found at mPFC-to-NAc, but not BLA-to-NAc, synapses (Figure 6). This is consistent with diet-induced plasticity in the mPFC (Dingess et al., 2017; Reichelt et al., 2019). We next expressed a Gi-coupled DREADD in the mPFC and activated these receptors located in mPFC terminals in the NAc by incubating slices from OPs in CNO (Figure 7A; ORs were not included because neither JF-Dep nor APV treatment altered their NAc CP-AMPAR transmission). We found that CNO treatment increased CP-AMPAR transmission compared to treatment of slices from the same rat with vehicle (Figure 7B). Control recordings performed by washing CNO onto slices with and without DREADD expression confirmed that DREADD activation results in a small, but significant decrease in AMPAR-mediated transmission. Thus, a relatively brief reduction in activity of mPFC-to-NAc inputs was sufficient to support synaptic “scaling-up” of CP-AMPARs.

Synaptic scaling in the NAc has been demonstrated using an NAc/mPFC neuronal co-culture system, however this involves increases in CI-AMPARs, not CP-AMPARs, induced by prolonged blockade (24-72h) of excitatory transmission by an AMPAR antagonist (Sun and Wolf, 2009; Werner et al., 2017). In hippocampal cultures, NMDAR blockade combined with action potential blockade is sufficient to recruit CP-AMPARs to synapses (Ju et al., 2004; Sutton et al., 2006), with some evidence that NMDAR blockade alone is sufficient (Sutton et al., 2006). To our knowledge, results here are the first demonstration of a similar acute effect in adult brain slices or within the NAc. Increases in CP-AMPAR transmission here occurred within two hours of NMDAR blockade, consistent with the scaling observed in culture (Turrigiano et al., 1998; Sutton et al., 2006; Sutton and Schuman, 2006). While this is a relatively rapid effect, the timescale is long enough that recruitment of existing receptors or the addition of newly synthesized receptors into synapse could contribute; these two possibilities are not mutually exclusive. However, in favor of the latter, CP-AMPAR recruitment in cultured hippocampal neurons requires protein synthesis (Sutton et al., 2006), as does the maintenance of CP-AMPARs at synapses in adult NAc slices (Scheyer et al., 2014; see below for further discussion).

The magnitude of APV-induced increases in CP-AMPAR transmission here was similar in OP-CH and OP-JF groups. This was unexpected, as we’ve previously found evidence for the extra-synaptic accumulation of CP-AMPARs in OPs maintained on junk-food (10d) vs chow. Thus, we predicted that the magnitude of CP-AMPAR increases would be greater in OP-JF vs OP-CH groups here. It is possible that NMDAR blockade resulted in a maximal recruitment of CP-AMPARs, masking group differences, or conversely that more time may be needed to observe group differences in synaptic CP-AMPAR recruitment. Regardless, the effect seen in the OP-CH group in light of the lack of recruitment seen in OR rats suggest inherent differences in the plasticity window between these populations. This is consistent with absence of diet-induced NAc AMPAR plasticity generally observed in OR rats (Oginsky et al., 2016; Ferrario, 2020).

We do not know why CP-AMPAR plasticity occurs more readily in OPs than ORs. To date, no evidence for basal differences in NAc CP-AMPAR synaptic transmission between these lines has been found (e.g., Figure 5,6; Oginsky et al., 2016), although floor effects could mask differences. However, basal NAc GluA1, but not GluA2, surface protein expression is lower in male OPs vs ORs (Derman and Ferrario, 2018b; Alonso-Caraballo et al., 2021) and basal intrinsic excitability of NAc medium spiny neurons is enhanced in OPs vs ORs (Oginsky et al., 2016; Oginsky and Ferrario, 2019). This combination may render OPs more sensitive to experience-induced plasticity. Finally, it’s worthwhile to note that JF-Dep also enhances NAc CP-AMPAR transmission in female OPs but not ORs (Nieto et al., 2023). However, this increase is transient, in contrast to persistent effects observed in males (Oginsky et al., 2016; Alonso-Caraballo et al., 2021). Thus it will be important to examine females in future.

### D. Conclusions

Overall, we find that a history of junk-food consumption and obesity-susceptibility interact to enhance food-motivated behavior and recruit NAc CP-AMPARs. These data provide further evidence that interactions between predisposition and diet-induced neurobehavioral plasticity likely contribute to weight gain and the maintenance of obesity. In light of modern diet culture, these data also emphasize the importance of understanding lasting changes that occur after stopping a sugary, fatty diet and set the stage for future studies linking these synaptic changes to behavioral outcomes. Finally, these data reveal novel insights into the mechanisms underlying CP-AMPAR recruitment in the NAc that involve synaptic scaling mechanisms. This has important implications for both cue-triggered food- and potentially drug-seeking behaviors.

## Acknowledgments

This work was supported in part by NIH grants R01DA044204, R01DK106188, R01DK115526, and R01DK130246 to CRF. TLF was supported by NIH grant T32DA007268. AMC was supported by NIH grant T32DA007281.

## Author Contributions

TLF designed and executed the experiments, analyzed data, and wrote this manuscript. AMC executed the experiments and wrote this manuscript. CRF designed experiments, analyzed data, and wrote this manuscript. All authors approved the final version of this work.

## Disclosures

The authors declare no competing financial interests.

## Notes

### Competing Interest Statement

The authors have declared no competing interest.

## REFERENCES

Alonso-Caraballo Y, Fetterly TL, Jorgensen ET, Nieto AM, Brown TE, Ferrario CR (2021) Sex specific effects of “junk-food” diet on calcium permeable AMPA receptors and silent synapses in the nucleus accumbens core. Neuropsychopharmacology 46:569–578.

Brown RM, Kupchik YM, Spencer S, Garcia-Keller C, Spanswick DC, Lawrence AJ, Simonds SE, Schwartz DJ, Jordan KA, Jhou TC, Kalivas PW (2017) Addiction-like Synaptic Impairments in Diet-Induced Obesity. Biol Psychiatry 81:797–806.

Cardinal RN, Parkinson JA, Hall J, Everitt BJ (2002) Emotion and motivation: the role of the amygdala, ventral striatum, and prefrontal cortex. Neurosci Biobehav Rev 26:321–352.

Christoffel DJ, Walsh JJ, Heifets BD, Hoerbelt P, Neuner S, Sun G, Ravikumar VK, Wu H, Halpern CH, Malenka RC (2021) Input-specific modulation of murine nucleus accumbens differentially regulates hedonic feeding. Nat Commun 12:2135.

Clem RL, Huganir RL (2010) Calcium-permeable AMPA receptor dynamics mediate fear memory erasure. Science 330:1108–1112.

Conrad KL, Tseng KY, Uejima JL, Reimers JM, Heng LJ, Shaham Y, Marinelli M, Wolf ME (2008) Formation of accumbens GluR2-lacking AMPA receptors mediates incubation of cocaine craving. Nature 454:118–121.

Counotte DS, Schiefer C, Shaham Y, O’Donnell P (2014) Time-dependent decreases in nucleus accumbens AMPA/NMDA ratio and incubation of sucrose craving in adolescent and adult rats. Psychopharmacology (Berl) 231:1675–1684.

Crombag HS, Sutton JM, Takamiya K, Holland PC, Gallagher M, Huganir RL (2008) A role for alpha-amino-3-hydroxy-5-methylisoxazole-4-propionic acid GluR1 phosphorylation in the modulatory effects of appetitive reward cues on goal-directed behavior. Eur J Neurosci 27:3284–3291.

Demos KE, Heatherton TF, Kelley WM (2012) Individual differences in nucleus accumbens activity to food and sexual images predict weight gain and sexual behavior. J Neurosci 32:5549–5552.

Derman RC, Ferrario CR (2018a) Junk-food enhances conditioned food cup approach to a previously established food cue, but does not alter cue potentiated feeding; implications for the effects of palatable diets on incentive motivation. Physiol Behav 192:145–157.

Derman RC, Ferrario CR (2018b) Enhanced incentive motivation in obesity-prone rats is mediated by NAc core CP-AMPARs. Neuropharmacology 131:326–336.

Derman RC, Bass CE, Ferrario CR (2020) Effects of hM4Di activation in CamKII basolateral amygdala neurons and CNO treatment on sensory-specific vs. general PIT: refining PIT circuits and considerations for using CNO. Psychopharmacology (Berl) 237:1249–1266.

Dingess PM, Darling RA, Derman RC, Wulff SS, Hunter ML, Ferrario CR, Brown TE (2017) Structural and Functional Plasticity within the Nucleus Accumbens and Prefrontal Cortex Associated with Time-Dependent Increases in Food Cue-Seeking Behavior. Neuropsychopharmacology 42:2354–2364.

Dong Y, Taylor JR, Wolf ME, Shaham Y (2017) Circuit and Synaptic Plasticity Mechanisms of Drug Relapse. J Neurosci 37:10867–10876.

Ferrario CR (2020) Why did I eat that? Contributions of individual differences in incentive motivation and nucleus accumbens plasticity to obesity. Physiol Behav 227:113114.

Ferrario CR, Labouèbe G, Liu S, Nieh EH, Routh VH, Xu S, O’Connor EC (2016) Homeostasis Meets Motivation in the Battle to Control Food Intake. J Neurosci 36:11469–11481.

Fetterly TL, Oginsky MF, Nieto AM, Alonso-Caraballo Y, Santana-Rodriguez Z, Ferrario CR (2021) Insulin Bidirectionally Alters NAc Glutamatergic Transmission: Interactions between Insulin Receptor Activation, Endogenous Opioids, and Glutamate Release. J Neurosci 41:2360–2372.

Fetterly TL, Basu A, Nabit BP, Awad E, Williford KM, Centanni SW, Matthews RT, Silberman Y, Winder DG (2019) α(2A)-Adrenergic Receptor Activation Decreases Parabrachial Nucleus Excitatory Drive onto BNST CRF Neurons and Reduces Their Activity In Vivo. J Neurosci 39:472–484.

Finger BC, Dinan TG, Cryan JF (2012) Diet-induced obesity blunts the behavioural effects of ghrelin: studies in a mouse-progressive ratio task. Psychopharmacology (Berl) 220:173–181.

Guo C, Ma YY (2021) Calcium Permeable-AMPA Receptors and Excitotoxicity in Neurological Disorders. Front Neural Circuits 15:711564.

Holland PC, Petrovich GD (2005) A neural systems analysis of the potentiation of feeding by conditioned stimuli. Physiol Behav 86:747–761.

Horstmann A, Busse FP, Mathar D, Müller K, Lepsien J, Schlögl H, Kabisch S, Kratzsch J, Neumann J, Stumvoll M, Villringer A, Pleger B (2011) Obesity-Related Differences between Women and Men in Brain Structure and Goal-Directed Behavior. Front Hum Neurosci 5:58.

Ju W, Morishita W, Tsui J, Gaietta G, Deerinck TJ, Adams SR, Garner CC, Tsien RY, Ellisman MH, Malenka RC (2004) Activity-dependent regulation of dendritic synthesis and trafficking of AMPA receptors. Nat Neurosci 7:244–253.

Levin BE, Dunn-Meynell AA, Balkan B, Keesey RE (1997) Selective breeding for diet-induced obesity and resistance in Sprague-Dawley rats. Am J Physiol 273:R725–730.

Man HY (2011) GluA2-lacking, calcium-permeable AMPA receptors–inducers of plasticity? Curr Opin Neurobiol 21:291–298.

Matikainen-Ankney BA, Legaria AA, Pan Y, Vachez YM, Murphy CA, Schaefer RF, McGrath QJ, Wang JG, Bluitt MN, Ankney KC, Norris AJ, Creed MC, Kravitz AV (2023) Nucleus Accumbens D(1) Receptor-Expressing Spiny Projection Neurons Control Food Motivation and Obesity. Biol Psychiatry 93:512–523.

Murdaugh DL, Cox JE, Cook EW, 3rd, Weller RE (2012) fMRI reactivity to high-calorie food pictures predicts short- and long-term outcome in a weight-loss program. Neuroimage 59:2709–2721.

Nieto AM, Catalfio AM, Papacostas Quintanilla H, Alonso-Caraballo Y, Ferrario CR (2023) Transient effects of junk food on NAc core MSN excitability and glutamatergic transmission in obesity-prone female rats. Obesity (Silver Spring) 31:434–445.

Oginsky MF, Ferrario CR (2019) Eating “junk food” has opposite effects on intrinsic excitability of nucleus accumbens core neurons in obesity-susceptible versus -resistant rats. J Neurophysiol 122:1264–1273.

Oginsky MF, Goforth PB, Nobile CW, Lopez-Santiago LF, Ferrario CR (2016) Eating ‘Junk-Food’ Produces Rapid and Long-Lasting Increases in NAc CP-AMPA Receptors: Implications for Enhanced Cue-Induced Motivation and Food Addiction. Neuropsychopharmacology 41:2977–2986.

Reichelt AC, Gibson GD, Abbott KN, Hare DJ (2019) A high-fat high-sugar diet in adolescent rats impairs social memory and alters chemical markers characteristic of atypical neuroplasticity and parvalbumin interneuron depletion in the medial prefrontal cortex. Food Funct 10:1985–1998.

Richardson NI and Roberts DC (1996) Progressive ratio schedules in drug self-administration studies in rats: a method to evaluate reinforcing efficacy, J. Neurosci. Methods 66: 1–11.

Robinson MJ, Burghardt PR, Patterson CM, Nobile CW, Akil H, Watson SJ, Berridge KC, Ferrario CR (2015) Individual Differences in Cue-Induced Motivation and Striatal Systems in Rats Susceptible to Diet-Induced Obesity. Neuropsychopharmacology 40:2113–2123.

Scheyer AF, Wolf ME, Tseng KY (2014) A protein synthesis-dependent mechanism sustains calcium-permeable AMPA receptor transmission in nucleus accumbens synapses during withdrawal from cocaine self-administration. J Neurosci 34:3095–3100.

Small DM (2009) Individual differences in the neurophysiology of reward and the obesity epidemic. Int J Obes (Lond) 33 Suppl 2:S44–48.

Stice E, Figlewicz DP, Gosnell BA, Levine AS, Pratt WE (2013) The contribution of brain reward circuits to the obesity epidemic. Neurosci Biobehav Rev 37:2047–2058.

Sun X, Wolf ME (2009) Nucleus accumbens neurons exhibit synaptic scaling that is occluded by repeated dopamine pre-exposure. Eur J Neurosci 30:539–550.

Sutton MA, Schuman EM (2006) Dendritic protein synthesis, synaptic plasticity, and memory. Cell 127:49–58.

Sutton MA, Ito HT, Cressy P, Kempf C, Woo JC, Schuman EM (2006) Miniature neurotransmission stabilizes synaptic function via tonic suppression of local dendritic protein synthesis. Cell 125:785–799.

Swinburn BA, Sacks G, Hall KD, McPherson K, Finegood DT, Moodie ML, Gortmaker SL (2011) The global obesity pandemic: shaped by global drivers and local environments. Lancet 378:804–814.

Tukey DS et al. (2013) Sucrose ingestion induces rapid AMPA receptor trafficking. J Neurosci 33:6123–6132.

Turrigiano GG, Leslie KR, Desai NS, Rutherford LC, Nelson SB (1998) Activity-dependent scaling of quantal amplitude in neocortical neurons. Nature 391:892–896.

Volkow ND, Wang GJ, Tomasi D, Baler RD (2013) Obesity and addiction: neurobiological overlaps. Obes Rev 14:2–18.

Werner CT, Murray CH, Reimers JM, Chauhan NM, Woo KK, Molla HM, Loweth JA, Wolf ME (2017) Trafficking of calcium-permeable and calcium-impermeable AMPA receptors in nucleus accumbens medium spiny neurons co-cultured with prefrontal cortex neurons. Neuropharmacology 116:224–232.

Wolf ME, Tseng KY (2012) Calcium-permeable AMPA receptors in the VTA and nucleus accumbens after cocaine exposure: when, how, and why? Front Mol Neurosci 5:72.

